# Dynamic protein phosphorylation in *Streptococcus pyogenes* during growth, stationary phase and starvation

**DOI:** 10.1101/2023.11.24.568605

**Authors:** Stefan Mikkat, Michael Kreutzer, Nadja Patenge

## Abstract

**Background:** Phosphorylation of proteins at serine, threonine, and tyrosine residues plays an important role in physiological processes of bacteria such as cell cycle, metabolism, virulence, dormancy, sporulation, and stationary phase functions. *Streptococcus pyogenes* possesses a single known transmembrane serine/threonine kinase belonging to the class of penicillin-binding protein and serine/threonine-associated (PASTA) kinases. To gain initial insights into the targets and dynamics of protein phosphorylation in *S. pyogenes* serotype M49 strain 591, we performed a proteomics and phosphoproteomics workflow using cultures from different growth conditions, stationary phase and starvation.

**Results:** The quantitative analysis of dynamic phosphorylation, which included a subset of 463 out of 815 identified phosphorylation sites, revealed two main types of phosphorylation events, distinguished by the growth phase in which they predominantly occur and their preference for either threonine or serine. A small group of phosphorylation events occurred almost exclusively at threonine residues of proteins related to the cell cycle and was enhanced in growing cells. Many of these phosphorylation sites are highly conserved targets of PASTA kinases in streptococci. The majority of phosphorylation events occurred during stationary phase or starvation, preferentially at serine residues. These phosphorylations may be important for regulatory processes in stationary phase or for persister cell formation, but their function and the kinases responsible for their formation need to be elucidated in further analyses. Moreover, our data indicate that the vast majority of proteins can be phosphorylated, but identification of their phosphopeptides depends on the sensitivity of the proteomic methods used.

**Conclusions:** PASTA kinase-dependent cell cycle regulation processes found in related bacteria are conserved in *S. pyogenes*, but account for only a small part of all phosphorylation events. Most phosphorylation events take place during stationary growth phase and starvation. This phenomenon has also been described for some other bacteria and may therefore be a general feature of bacterial protein phosphorylation.

## 1 Introduction

Protein phosphorylation is one of the most important and best studied post-translational modifications involved in the regulation of various biological processes (Macek et al., 2019). Phosphorylation alters the physicochemical properties of a protein and can thereby control cellular processes, for example by inducing conformational changes in the active site of the protein or regulating protein-protein interactions (Pereira et al., 2011). The opposing action of phosphorylating kinases and dephosphorylating phosphatases enables rapid and precise regulation.

In bacteria, signal transduction by two-component systems involving phosphorylation of histidine and arginine residues has traditionally been considered the almost exclusive form of protein phosphorylation. However, during the last two decades it has become increasingly evident that phosphorylation at serine, threonine and tyrosine residues plays vital roles in several physiological processes, such as cell cycle and cell wall synthesis, bacterial metabolism, virulence, dormancy, sporulation and stationary phase functions (Macek et al., 2019).

Enrichment of phosphopeptides using immobilized metal affinity chromatography (IMAC) or TiO_2_ affinity chromatography (Thingholm et al., 2009) with subsequent LC-MS/MS analysis has been established as the method of choice for site-specific serine/threonine/tyrosine bacterial phosphoproteomics. Early pioneering projects using this approach typically identified near one hundred phosphorylation sites in the phosphoproteomes of *Bacillus subtilis* (Macek et al., 2007), *E. coli* (Macek et al., 2008), *Lactococcus lactis* (Soufi et al., 2008) and other bacteria (reviewed by Yagüe et al., 2019). This number increased dramatically to several thousand with technological advances in qualitative and quantitative mass spectrometry, data management and phosphopeptide sample preparation (Lin et al., 2018; Prust et al., 2021; Garcia-Garcia et al., 2022). Despite these advances, there are major gaps in our knowledge of the functions of protein phosphorylation in bacteria. The kinases/phosphatases that regulate the hundreds and thousands of identified phosphorylation events, as well as the full range of native substrates of known kinases/phosphatases, are largely unknown. Functional redundancy of different kinases and/or substrate promiscuity complicate their elucidation (Pereira et al., 2011). In addition, there are no unique sequence motifs of bacterial kinase targets.

Bacterial phosphorylation at serine and threonine residues is mainly catalyzed by eukaryotic-like Ser/Thr kinases (eSTKs) that possess a catalytic domain sharing a common fold with eukaryotic protein kinases. Ubiquitous in Actinobacteria and Firmicutes are transmembrane eSTKs composed of a cytoplasmic catalytic kinase domain linked by a transmembrane segment to an extracellular domain consisting of a variable number of PASTA (penicillin-binding protein and serine/threonine-associated) repeats (Pereira et al., 2011; Manuse et al., 2016). Binding of ligands such as beta-lactam compounds and cell wall fragments to the PASTA domains triggers the kinase activity (Yeats et al., 2002). PASTA kinase activity affected virulence of streptococci and other firmicutes in animal models (reviewed by Kohler et al., 2020)

Tyrosine phosphorylation is performed by bacterial idiosyncratic tyrosine kinases (BY-kinases) (Grangeasse et al., 2007; Mijakovic et al., 2016) and by the family of ubiquitous bacterial kinases (UbK) (Nguyen et al., 2017). In addition, atypical and novel kinases contribute to the diversity of bacterial systems for phosphorylation of serine, threonine and tyrosine (Castro-Roa et al., 2013; Mijakovic et al., 2016; Rajagopalan and Dworkin, 2020).

*Streptococcus pyogenes* is a host-adapted human pathogen responsible for a wide range of clinical manifestations, including asymptomatic infection, superficial self-limiting infections, invasive life-threatening diseases, and autoimmune sequelae. Together, the global burden of disease caused by *S. pyogenes* is high (Carapetis et al., 2005). The course of an infection depends on the appropriate expression of specific virulence factors and on the successful adaption to the environment within the host (Brouwer et al., 2023). Control of virulence gene expression by stand-alone transcription factors and two-component systems are known to play a role in virulence determination in *S. pyogenes* (Kreikemeyer et al., 2003; Vega et al., 2022). An additional level of bacterial gene expression control is provided by small regulatory RNAs (Tesorero et al., 2013; Le Rhun et al., 2016). Furthermore, protein phosphorylation could play a crucial role in the pathogenicity of *S. pyogenes*, as already known for other bacteria (Kohler et al., 2020).

*S. pyogenes* possesses one PASTA kinase and its cognate protein phosphatase, which are products of cotranscribed genes. In *S. pyogenes* M1, the kinase SP-STK was autophosphorylated *in vitro* on threonine residues and was dephosphorylated by the phosphatase SP-STP. Reversible phosphorylation by SP-STK/SP-STP affected cell division/septation, shape, size, expression and surface display of certain proteins, and interaction of *S. pyogenes* with eukaryotic cells. While removal of SP-STK or its kinase or PASTA domain altered growth and morphological characteristics differently, SP-STP proved to be an essential enzyme in *S. pyogenes* M1 (Jin and Pancholi, 2006). Hereafter, the PASTA kinase (Spy49_1257c) and the cognate phosphatase (Spy49_1259c) of *S. pyogenes* M49 will be referred to as SP-STK and SP-STP, respectively, consistent with the proteins from *S. pyogenes* M1 (Jin and Pancholi, 2006).

*S. pyogenes* has no BY-kinase, but possesses a ubiquitous bacterial kinase (Ubk) (Nguyen et al., 2017; Kant and Pancholi, 2021) and a low molecular weight protein tyrosine phosphatase (Kant et al., 2015). Recently, the first phosphoproteome dataset of *S. pyogenes* was published (Birk et al., 2021). However, compared to *Streptococcus pneumoniae* and other streptococci, little is known about the role of protein phosphorylation in *S. pyogenes*. In this work, we cultivated *S. pyogenes* M49 in different media to investigate the dynamics of protein phosphorylation during growth, stationary phase and starvation.

## 2 Materials and Methods

### 2.1 Bacterial culture conditions

*S. pyogenes* serotype M49 strain 591 (Kaufhold et al., 1992; Patenge et al., 2021) was cultured in Todd-Hewitt broth supplemented with 0.5% yeast extract (THY; Oxoid, Thermo Fisher Scientific, Darmstadt, Germany) at 37°C under a 5% CO_2_/20% O_2_ atmosphere. Overnight cultures were diluted 1:20 in 40 ml THY and grown to an OD_600_ = 0.8. For media exchange, bacteria were centrifuged at 4,000 g for 20 min. The pellet was suspended in either THY or chemically defined medium (van de Rijn and Kessler, 1980) without carbon source (CDM-) or with 1% fructose (CDMF). Cells were harvested at different time points as indicated (Fig. 1, Additional file X: Fig. S1). The culture was centrifuged at 4,000 g for 10 min at 4°C. The pellet was washed in cold PBS, subsequently shock frozen in liquid nitrogen, and stored at -80°C.

### 2.2 Sample preparation for proteomics

Ice-cooled bacterial cells were disrupted with glass beads using Precellys 24 homogenizer (peqLab Biotechnologie GmbH, Erlangen, Germany) in non-denaturing buffer containing 10 mM Tris/HCl, pH 7.4, 138 mM NaCl, 2.7 mM KCl, 1 mM MgCl_2_. Immediately thereafter, Tris-HCl, pH 8.0 and sodium deoxycholate (SDC) were added from stock solutions to obtain final concentrations of 50 mM Tris-HCl and 2% SDC. Samples were incubated at 95 °C for 5 min before the protein extracts were aspirated from the glass beads and further sonicated for 10 min using a bath sonicator. The raw cell extract containing cell debris was used for the subsequent sample processing steps including proteolytic digestion. Protein concentration was measured using the Bio-Rad protein assay (Bio-Rad, Munich, Germany).

Reduction/alkylation was performed for 15 min at 37 °C after addition of 1/10 volume reduction/alkylation reagent containing 100 mM tris(2-carboxyethyl)phosphine hydrochloride (TCEP) and 400 mM 2-chloroacetamide (CAM) (Humphrey et al., 2018). Next, methanol/chloroform precipitation was performed as described by Potel et al. (2018). The precipitate was dissolved in digestion buffer composed of 100 mM Tris-HCl, pH 8.0, 1% SDC, 5 mM TCEP, and 20 mM CAM. Sequencing grade trypsin (Promega GmbH, Walldorf, Germany) was added to obtain an enzyme/protein ratio of approximately 1:100 and digestion was performed at 37°C for about 16 h. Afterwards cell debris was removed by centrifugation. The supernatant was acidified to a final concentration of 0.7% trifluoroacetic acid (TFA), mixed vigorously, and the precipitated SDC was pelleted by centrifugation at 13,000 rpm for 10 min. Finally, the peptide solutions were desalted with OASIS HLB 1cc 30 mg Vac Cartridges (Waters, Manchester, UK) and the eluate was fivefold concentrated using a centrifugal evaporator. Peptide concentrations were measured using the Invitrogen Qubit protein assay kit (Thermo Fisher Scientific, Darmstadt, Germany).

### 2.3 Phosphopeptide Enrichment

Peptide amounts of 100 µg or 250 µg were evaporated to dryness using a centrifugal evaporator. MagReSyn TiO_2_ and MagReSyn Ti-IMAC hyperporous magnetic microparticles (ReSyn Biosciences, Edenvale, Gauteng, South Africa) were used as a mixture to take advantage of combined enrichment chemistries (Tape et al., 2014). Compositions of loading buffer (1 M glycolic acid in 80% acetonitrile (ACN) and 5% TFA), wash buffer 1 (80% ACN, 1% TFA), wash buffer 2 (10% ACN, 0.2% TFA), and elution buffer (1% NH_4_OH) corresponded to the manufacturer’s recommendation for use with MagReSyn TiO_2_ microparticles. For the enrichment of phosphopeptides from one sample, 15 µl Ti-IMAC and 7.5 µl TiO_2_ were mixed and equilibrated with loading buffer.

Dried peptides were dissolved in 200 µl loading buffer, freed from insoluble material by centrifugation and transferred to the pellet of equilibrated microparticles. The phosphopeptides were bound to the microparticles during an incubation of 20 min at room temperature with constant mixing. Three consecutive washes of two minutes each were then performed with 100 µl loading buffer, wash buffer 1, and wash buffer 2. Bound phosphopeptides were eluted from the microparticles with 80 µl of elution buffer with gentle mixing for 10 minutes. The buffer containing the eluted peptides was transferred to a protein LoBind tube (Eppendorf, Hamburg, Germany) containing 20 µl of 10% formic acid (FA). The elution was repeated with 80 µl of elution buffer for 5 min, and both eluates were pooled and frozen. Subsequently, the samples were evaporated to near dryness using a centrifugal evaporator. Desalting of the phosphopeptides was performed on StageTips containing one C18 disk as described (Rappsilber et al., 2007). Finally, phosphopeptides were dissolved in 20 µl of 2% ACN, 0.1% FA.

### 2.4 Mass spectrometry

LC-MS analyses were carried out using a nanoAcquity UPLC system (Waters, Manchester, UK) coupled to a Waters Synapt G2-S mass spectrometer via a NanoLockSpray ion source as described (Klähn et al., 2021). Mobile phase A contained 0.1% FA in water and mobile phase B contained 0.1% FA in acetonitrile. Peptides were trapped and desalted using a precolumn (ACQUITY UPLC Symmetry C18, 5 µm, 180 µm x 20 mm, Waters) and subsequently separated on an analytical column (ACQUITY UPLC HSS T3, 1.8 µm, 75 µm x 250 mm, Waters) at a flow rate of 300 nl/min using a gradient from 3% to 32% B over 90 min. For the analysis of the total proteome, 70 ng of peptides according to the results of the Invitrogen Qubit protein assay (Thermo Fisher Scientific, Darmstadt, Germany) supplemented with 40 fmol of Hi3 Phos B standard for protein absolute quantification (Waters) were injected. For analyses of phosphopeptides, typically 10% of the enriched sample was used.

For both, total proteome and phosphoproteome measurements, the Synapt G2-S instrument was operated in data-independent mode with ion-mobility separation as an additional dimension of separation (referred to as HDMS^E^). By executing alternate scans at low and elevated collision energy of each 0.6 sec, information on precursor and fragment ions, respectively, was acquired. (Distler et al., 2014; Klähn et al., 2021). Either duplicate or triplicate measurements were performed, depending on the experiment. Measurements of phosphoproteomes were additionally conducted in data-dependent mode (Fast DDA) of the Synapt G2-S mass spectrometer. Typical parameters used were as follows. Following MS survey scans of 0.2 s, the instrument was switching to MS/MS acquisition if the intensity of individual ions exceeded the threshold of 50,000 counts per second. Up to three ions which charge states between 2+ and 4+ were selected from a single MS survey scan. MS/MS scan rate was set to 0.2 s. The instrument returned to MS survey scan if the TIC threshold of 600,000 counts was reached or after 2.1 s of MS/MS acquisition. Dynamic exclusion of already selected precursor ions was set to 12 s. In addition, inclusion lists generated from HDMS^E^ analyses of the respective experiments were used for some DDA acquisitions. Here, exclusively ions from the inclusion list were selected for MS/MS provided that the precursor ion threshold exceeded 8,000 counts per second and mass and retention time of an ion were within 40 mDa and 90 s windows, respectively, of entries on the inclusion list.

### 2.5 Data processing, protein identification and quantification

#### 2.5.1 Total proteome

Progenesis QI for Proteomics version 4.1 (Nonlinear Dynamics, Newcastle upon Tyne, UK) was used for raw data processing, protein identification, and label-free quantification as described (Klähn et al., 2021). Proteins were identified using a database of 1,701 protein sequences from *Streptococcus pyogenes* serotype M49, strain NZ131 (UniProt release 2021_02) appended with the sequences of rabbit phosphorylase B (P00489) and porcine trypsin. Proteins were quantified by the absolute quantification Hi3 method using Hi3 Phos B Standard (Waters) as reference (Silva et al., 2006). Results were given as fmol on column.

#### 2.5.2 Phosphoproteome

Raw data from data independent (HDMS^E^) acquisitions were processed with Progenesis QI for Proteomics. Then, three different strategies for peptide and protein identification were applied to the same data: (i) ion accounting search, (ii) Mascot search, and (iii) spectral library search.

For identification using the ion accounting algorithm implemented in Progenesis, a database containing 1,701 protein sequences from *S. pyogenes* serotype M49, strain NZ131 (UniProt release 2021_02) was used. Two missing cleavage sites were allowed, carbamidomethylation of cysteine residues was set as fixed modification and methionine oxidation, asparagine deamidation, phosphorylation of serine, threonine and tyrosine residues, and phosphoglyceryl modification of lysine residues were considered as variable modification. The false discovery rate was set to 1%. Peptides were required to be identified by at least five fragment ions. Subsequently peptide ion data were filtered to retain only peptide ions that met the following criteria: (i) identified in at least two samples within the dataset, (ii) minimum ion score of 5.5, (iii) mass error below 13.0 ppm, and (iiii) at least 8 amino acid residues in length. Identifications based on charge state deconvolution were removed. Deamidation of asparagine was only accepted, if the asparagine residue was followed by glycine (Mikkat et al., 2013). Subsequently, the fragment spectra of all remaining phosphopeptides were checked manually for plausibility of identification. Questionable peptide identifications were removed while reliable peptides with ambiguously assigned phosphorylation sites were tagged for further analyses.

To identify HDMS^E^ data with the Mascot search engine, peak lists in Mascot Generic Format (MGF files) were generated in Progenesis. The number of exported fragment ions per spectrum was limited to 80 and deisotoping and charge deconvolution was enabled. The MGF files were searched against the *S. pyogenes* database using Mascot (version 2.6.2) applying the same enzyme specificity, fixed and variable modifications used for identification with the ion accounting algorithm. Peptide mass tolerance was set to 13 ppm and fragment mass tolerance to 0.02 Da. The significance threshold was adjusted to p < 0.01. Resulting protein false positive rates were between zero and 0.44% for the different experiments. The results data were imported into Progenesis and mapped to the peptide features. Then, phosphopeptides identified in only one sample of the data set were deleted. Deamidation of asparagine was only accepted, if the asparagine residue was followed by glycine. The confidence of phosphorylation site localization was assessed using the Mascot Delta Score (Savitski et al., 2011). The maximum and mean Delta Score was calculated for multiple mass spectra belonging to the same feature. The maximum delta score was used to generate a list of all phosphopeptides, while the mean delta score was used to select phosphopeptides for quantitative analysis.

To search HDMS^E^ data against a spectral library, spectral libraries were created in Progenesis with phosphopeptides that were unambiguously identified by DDA of the respective samples. Spectral library searches were performed using a peptide mass tolerance of 13 ppm, a fragment mass tolerance of 20 ppm, and a retention time window of 0.5 min. At least five matching fragments per peptide were required. The ion accounting, Mascot, and spectral library search results of the HDMS^E^ data were further processed using Microsoft Office.

Raw data from data dependent (DDA) acquisitions were also processed with Progenesis QI for Proteomics. Export of peak lists, peptide identification using the Mascot search engine and further processing of search results were as described for the HDMS^E^ data. Hierarchical clustering was performed using the Interactive CHM Builder (Ryan et al., 2022) at https://build.ngchm.net/NGCHM-web-builder/. The mass spectrometry proteomics data have been deposited to the ProteomeXchange Consortium via the PRIDE (Perez-Riverol et al., 2022) partner repository with the dataset identifier PXD044423 and 10.6019/PXD044423.

## 3 Results

### 3.1 Experimental rationale

This study was designed to provide initial insights into targets and dynamics of protein phosphorylation in *S. pyogenes*. For this purpose, bacteria were first grown to late exponential phase in THY broth, then transferred to three different culture media and harvested at different time points up to 24 hours. We used rich THY broth containing 0.2% glucose providing optimal growth conditions, chemically defined medium containing 1% fructose (CDMF) to induce growth on a single carbon source, and chemically defined medium without a carbon source (CDM-) to provoke starvation. In the first experiment, we found that the number of phosphorylation sites and the overall degree of protein phosphorylation increased sharply in the stationary growth phase and under starvation conditions. Therefore, the cultivation time was extended to 72 hours in the second experiment with an otherwise identical experimental setup (Fig. 1). A third experiment utilizing only THY broth was designed to find out whether the phosphorylation pattern changed within 40 min after the bacteria were transferred from extended stationary phase to fresh medium ( Supplementary Figure S1). In total, proteomes and phosphoproteomes of 29 bacterial cultures were included in this study.

### 3.2 The proteome during growth in different culture media

Between 930 und 972 proteins identified by at least two unique peptides were quantified in the three experiments. 872 proteins were common to all experiments, while in total 1038 proteins were found, corresponding to 61% of the predicted proteins of *S. pyogenes*. The numbers of common and total proteins increased to 936 and 1121, respectively, when proteins whose identification was based on a single peptide were included (Supplementary Table S1, Supplementary Figure S2). Quantitative data of the proteome were collected primarily to allow normalization of phosphopeptide amounts with the corresponding protein amounts. Therefore, the influence of culture conditions on the proteome will only be shown using a few examples.

The levels of many proteins were increased in the stationary growth phase in THY. For example, nine proteins of the histidine degradation pathway whose genes are located in a common chromosomal region (spy49_1724 to spy49_1731, as well as spy49_1723c on the complementary strand) were significantly upregulated only in stationary THY cultures (Supplementary Figure S3). The same applies to the nine subunits of the V-type ATP synthase encoded by the contiguous genes spy49_0129 to spy49_0137 (Supplementary Figure S4). We hypothesize that the V-type ATPase is involved in pH regulation of cells grown in THY. Growth on fructose induced or enhanced the expression of several proteins, most strikingly of a group of nine proteins with less characterized functions encoded by the consecutive genes Spy49_0450 to Spy49_0460 (Supplementary Figure S5). With regard to the phosphoproteomic analysis, it is important to point out that the expression profiles of these and other proteins indicate differences in the time course of metabolic adaptation to fructose utilization between experiments 1 and 2.

There were also various proteins whose abundance decreased in the stationary phase. These were often surface-exposed. Cultivation in CDM- did not allow growth of the bacterial cultures, as illustrated by the continuous decrease in optical density (Figure 1). Nevertheless, to standardize sample designations with the other culture conditions, we have also used the terms exp, stat, and late stat for the CDM- cultures in the tables and figures. Cultivation in CDM- caused lesser effects on the proteome, mainly a decrease in surface-exposed and membrane proteins with increasing duration of cultivation. However, many ribosomal proteins, such as L28 and L35, were more reduced in CDM- than in the other media (Supplementary Figure S6). Despite the considerable differences in the expression of many proteins, about 60% of the proteins remained almost unaffected (fold change < 2) by the culture conditions studied.

**Figure 1.**
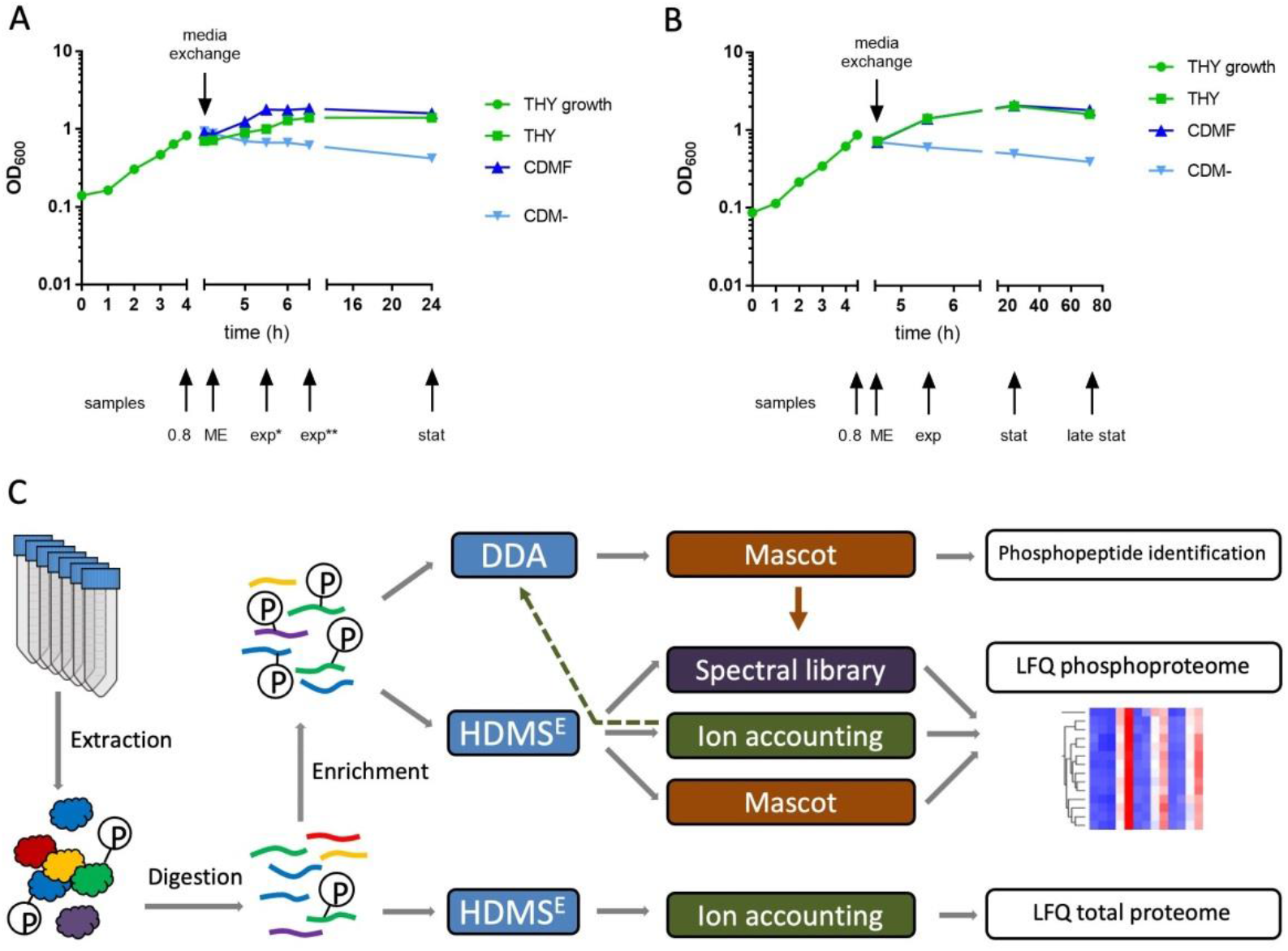
Bacterial growth and proteomics workflow. *S. pyogenes* was grown in 40 ml THY to an OD600 = 0.8 (green circles). For media exchange (ME), bacterial cultures were centrifuged and pellets were suspended in either THY (THY, green squares), CDM without carbon source (CDM-, light blue triangles) or CDM with 1% fructose (CDMF, dark blue triangles). Sample collection is indicated by arrows. exp: exponential growth phase (one doubling), stat: stationary phase, late stat: late stationary phase. **(A)** In the first experiment, *S. pyogenes* was cultured for 24 hours. **(B)** In the second experiment, *S. pyogenes* was cultured for 72 hours. **(C)** Experimental workflow for proteomic and phosphoproteomic nano LC-MS/MS analysis using a Synapt G2-S mass spectrometer and the analysis software Progenesis QI for proteomics. For label-free quantification of the total proteome, tryptic digests of the extracted proteins were subjected to data-independent HDMS^E^ acquisition. The ion accounting algorithm implemented in Progenesis was used for peptide and protein identification. Phosphopeptides were enriched using a mixture of TiO_2_ and Ti-IMAC hyperporous magnetic microparticles and subjected to both data-dependent (DDA) and HDMS^E^ acquisition. Peak lists from the DDA measurements were exported to Mascot for identification. The HDMS^E^ data were subjected to peptide identification by Mascot, ion accounting and comparison with a spectral library assembled from phosphopeptides identified in the DDA/Mascot approach (indicated by the brown arrow). For some DDA acquisitions, precursor selection was based on an inclusion list generated from the results of the HDMS^E^/ion accounting approach (indicated by the dashed arrow).

### 3.3 Identification of phosphopeptides by different search strategies

To study dynamic protein phosphorylation, we used label-free, data-independent LC-HDMS^E^ acquisition, which collects high-quality MS data across the entire chromatographic peak width (Silva et al., 2006). Using Progenesis QI for proteomics (Waters) for data analysis, this approach provides reliable quantification of unmodified peptides and proteins (Pappesch et al., 2017; Klähn et al., 2021), but the proprietary ion accounting algorithm for peptide and protein identification was found to be error-prone in identifying phosphopeptides. Increasing the stringency of the ion accounting search enabled confident identification of the peptide sequences, but the location of the phosphorylation site was often ambiguous and required time-consuming manual validation. As an alternative to the ion accounting algorithm, peak lists were searched using the Mascot search engine. This strategy provided reliable phosphopeptide identifications supported by a Mascot Delta Score-based (MD-score) phosphosite localization probability (Savitski et al., 2011). Additionally, we used spectral libraries assembled from high-confidence MS/MS spectra of phosphopeptides obtained from the same experiments by data-dependent analysis (DDA) and Mascot search. Thus, four data sets were generated from each of the three experiments: (i) HDMS^E^ data searched with the ion accounting algorithm, (ii) HDMS^E^ data searched with Mascot, (iii) HDMS^E^ data searched against a spectral library, and (iiii) DDA data searched with Mascot (Fig. 1). For each experiment, the three identification results of the HDMS^E^ data were combined in a single Excel sheet, whereas the DDA results are shown separately (Supplementary Table S2). Identification of the same feature using either ion accounting, Mascot or a spectral library search always resulted in concurring peptide sequences, but sometimes differing phosphorylation site localizations which required further critical revision.

Here we first consider the phosphopeptides without taking into account the phosphorylation site (Table 1). Most phosphorylated peptides were found in the second experiment, which included the largest number of different growth conditions. In all experiments, a total of 352 phosphoproteins derived from 955 peptide sequences were identified. The concordance between the identification of phosphorylated peptide sequences in the three experiments is shown in Figure 2A and 2B for the HDMS^E^ and DDA measurements, respectively. Almost all peptides identified in experiment 1 were also found in experiment 2, which additionally included samples from the late stationary phase. Experiment 3, which also contained late stationary phase cultures, yielded comparable numbers to experiment 2. This indicates that the number of phosphorylation sites increased during the prolonged stationary phase. The lower number of identifications in experiment 3 by DDA (Figure 2B) is related to the fact that a smaller number of measurements were performed.

**Figure 2.**
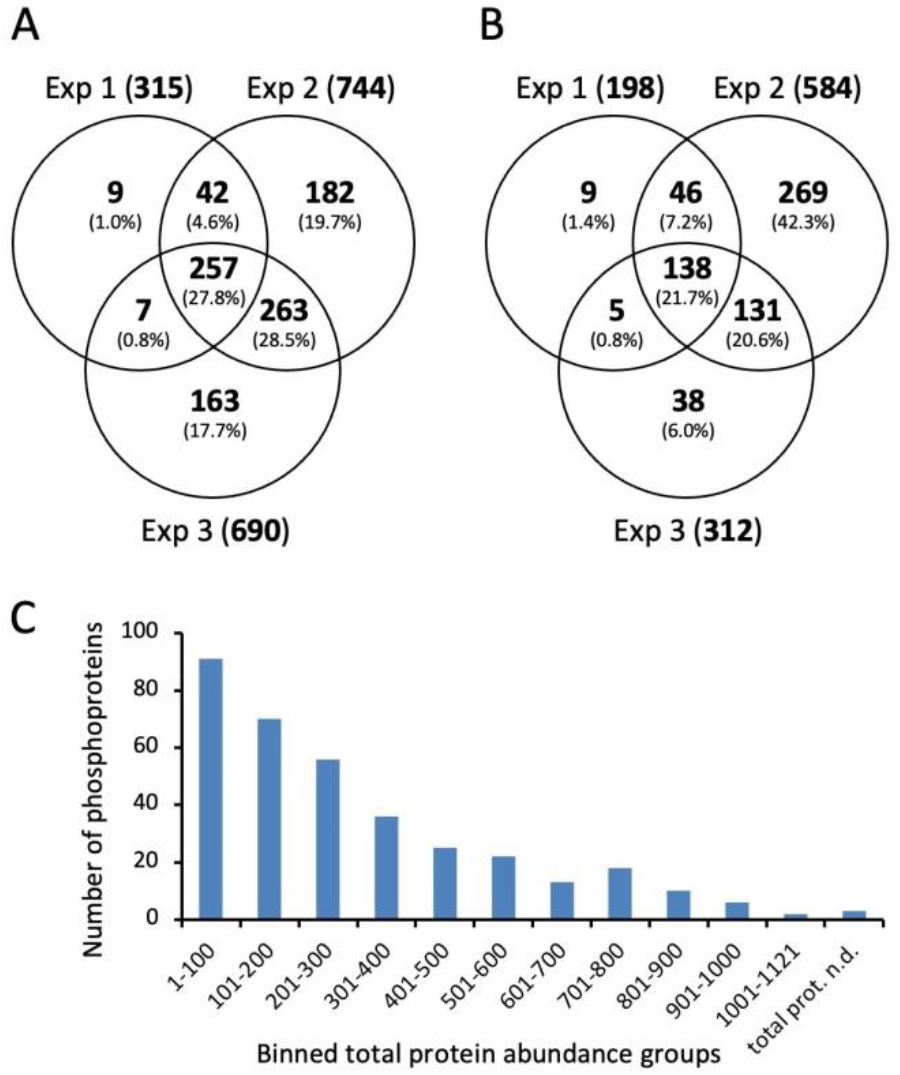
Identification of phosphorylated peptides without considering the phosphorylation site. **(A)** Venn diagram showing numbers and percentages of phosphorylated peptides identified in three experiments after HDMS^E^ acquisition, **(B)** Venn diagram showing numbers and percentages of phosphorylated peptides identified in three experiments after DDA acquisition, **(C)** Dependency of phosphopeptide identification on protein abundance. Each bar represents 100 proteins ordered according to decreasing abundance.

**Table 1.**
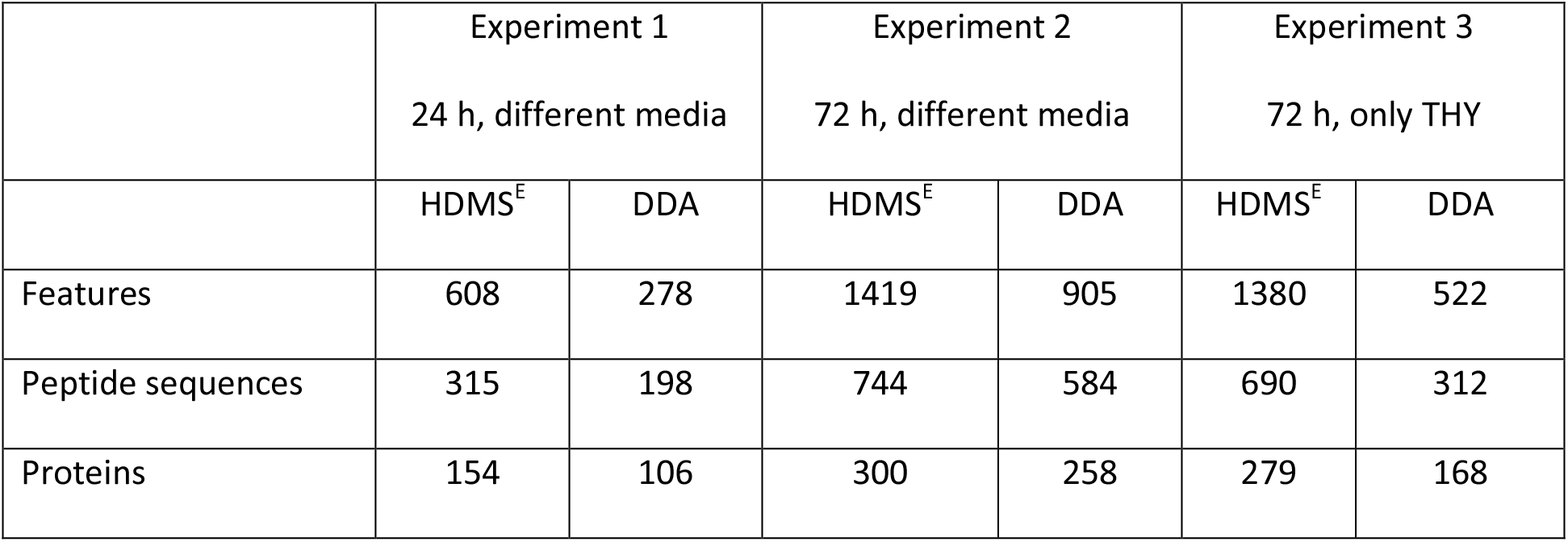
Number of phosphopeptide features, phosphorylated peptides without taking into account the phosphorylation site, and phosphoproteins identified in three independent experiments.

|  | Experiment 1 |  | Experiment 2 |  | Experiment 3 |  |
| --- | --- | --- | --- | --- | --- | --- |
|  | 24 h, different media |  | 72 h, different media |  | 72 h, only THY |  |
|  | HDMS<sup>E</sup> | DDA | HDMS<sup>E</sup> | DDA | HDMS<sup>E</sup> | DDA |
| Features | 608 | 278 | 1419 | 905 | 1380 | 522 |
| Peptide sequences | 315 | 198 | 744 | 584 | 690 | 312 |
| Proteins | 154 | 106 | 300 | 258 | 279 | 168 |

There was a strong correlation between the protein abundance determined in proteomic analysis and the identification of phosphopeptides (Figure 2C). We identified phosphorylated forms of 91 of the one hundred most abundant proteins. Of the nine nonphosphorylated proteins, three were surface-exposed proteins, such as metal ABC transporter substrate-binding lipoprotein mtsA and foldase protein PrsA whose phosphorylation by cytoplasmic kinases would not be expected.

### 3.4 Creating a list of phosphorylation sites

To generate a list of high-confidence phosphorylation sites, features identified in the Mascot search with a localization probability below 75% were removed independently from the results of the other search strategies. We applied the threshold of 75% but manually validated the localization of phosphorylation sites with a localization probability lower than 95% to avoid the exclusion of informative phosphorylation sites, e.g. from the PASTA kinase SP-STK, from the analysis. Ion accounting results with uncertain site localization were retained only if the site matched confident site localization from the Mascot search for the same feature. Initially, a promising strategy seemed to be to assemble a spectral library of all reliably identified phosphopeptides from all experiments to facilitate identification. Unfortunately, the positional variants of phosphopeptides were not reliably distinguished when searching the spectral library. Therefore, results from spectral library searches were used only if they were confirmed by the Mascot search.

Finally, the verified phosphopeptides from the HDMS^E^ and DDA approaches of the three experiments were merged and aligned to 15 amino acid stretches with a central phosphorylation site. Only phosphorylation sites identified in at least two conditions, e.g. from the same peptide ion identified in different experiments or with different search strategies, or from different peptide ions containing the same phosphorylation site, were included in the final list. Using these stringent criteria, a list of 815 phosphorylation sites was created (Supplementary Table S3). 240 sites (29.4%) were identified in all three experiments and 592 sites (72.6%) were identified in at least two experiments despite the different designs of the three experiments (Supplementary Figure S7A). As might be expected, the different data acquisition and search strategies complement each other. However, most phosphorylation sites (78.5%) were identified by searching the HDMS^E^ data using Mascot. (Supplementary Figure S7B).

The proportion of phosphorylated serine, threonine, and tyrosine residues was 73%, 21%, and 6%, respectively. The 815 phophorylation sites were distributed among 294 proteins. Of these, ten proteins contained at least ten phosphorylation sites; first among these was elongation factor Tu (EF-Tu) with 20 sites. On the other hand, a single phosphorylation site was found in 131 proteins (Supplementary Table S4).

With few exceptions, the identified phosphopeptides were phosphorylated at one single site. However, we observed many positional isomers of singly phosphorylated peptides with reproducible differences in chromatographic retention times as exemplified in the supplemental material for five phosphorylation sites on the same tryptic peptide of the general stress protein spy49_1001c (Supplementary Figure S8).

### 3.5 Phosphopeptide enrichment specificity

To evaluate the specificity of phosphopeptide enrichment and the efficiency of identification, we inspected in Progenesis the plot of m/z versus retention time combining all HDMS^E^ data of the second experiment (Supplementary Figure S9). The most abundant peptide ions derived from one large non-phosphorylated peptide (MYEQAAAAQQAAQGAEGAQANDSANNDDVVDGEFTEK) of chaperone protein DnaK. Non-phosphorylated peptides of the chaperonin GroEL were also highly abundant. Among phosphopeptides, prominent ion signals derived from the PASTA kinase SP-STK and cell-division initiation protein (DivIVA). However, many abundant signals were not identified including a striking pattern of ions with decreasing retention time at increasing m/z. It comprises a series of seven triply charged peptides, each differing by a mass difference of 162 Da, all of which could be assigned to the C-terminal peptide of the foldase protein PrsA (PrtM1) by manual analyses of fragment spectra. A mass increase of 162 Da indicates a sugar moiety which was also confirmed by an error tolerant Mascot search (Creasy and Cottrell, 2002). However, the mass spectra did not provide further information about the exact location and nature of the modification (Supplementary Figure S10). Foldase PrsA is a membrane-bound lipoprotein with peptidyl-prolyl cis-trans isomerase activity that supports folding of exported proteins and contributes to bacterial virulence (Alonzo III et al., 2011; Lin et al., 2018). Its identification as a putative glycoprotein may prompt further studies that could provide new insights into the virulence mechanisms of *S. pyogenes*.

Surprisingly, the error-tolerant Mascot search revealed frequently occurring lysine phosphoglycerylations (PGK) among the enriched phosphopeptides. This modification has been described for bacteria (Boel et al., 2004) and mammals (Moellering and Cravatt, 2013), but has not been studied further to date. We identified several hundred PGK-modified peptides. Considering the structural and functional differences between S/T/Y phosphorylation and phosphoglycerylation, we will publish the results separately. Specificity of enrichment was assessed by both the number of unique peptides and the molar ratios of the second experiment. Out of a total of 3581 peptides identified, the proportion of S/T/Y-phosphorylated peptides was 26%, while 63% were not phosphopeptides and 11% of peptides were modified by lysine phosphoglycerylation. Label-free quantification revealed percentages of S/T/Y-phosphorylated peptides, nonphosphorylated peptides, and PGK-modified peptides of 24%, 73%, and 3%, respectively.

### 3.6 Quantitative analysis of dynamic protein phosphorylation

For quantitative analysis of dynamic protein phosphorylation, we considered a subset of 463 phosphorylation sites quantified in HDMS^E^ analyses from at least two of the three experiments (Figure 3A). For each phosphorylation site, the amounts of all associated phosphopeptide ions, i.e., species with different charge states, missed cleavages, and additional modifications, were summed (Supplementary Table S5). First, we analyzed the quantitative distribution of phosphorylation events at serine, threonine, and tyrosine over the course of the experiments. Because of its dominance, threonine phosphorylation of SP-STK was quantified separately (Figure 3).

**Figure 3.**
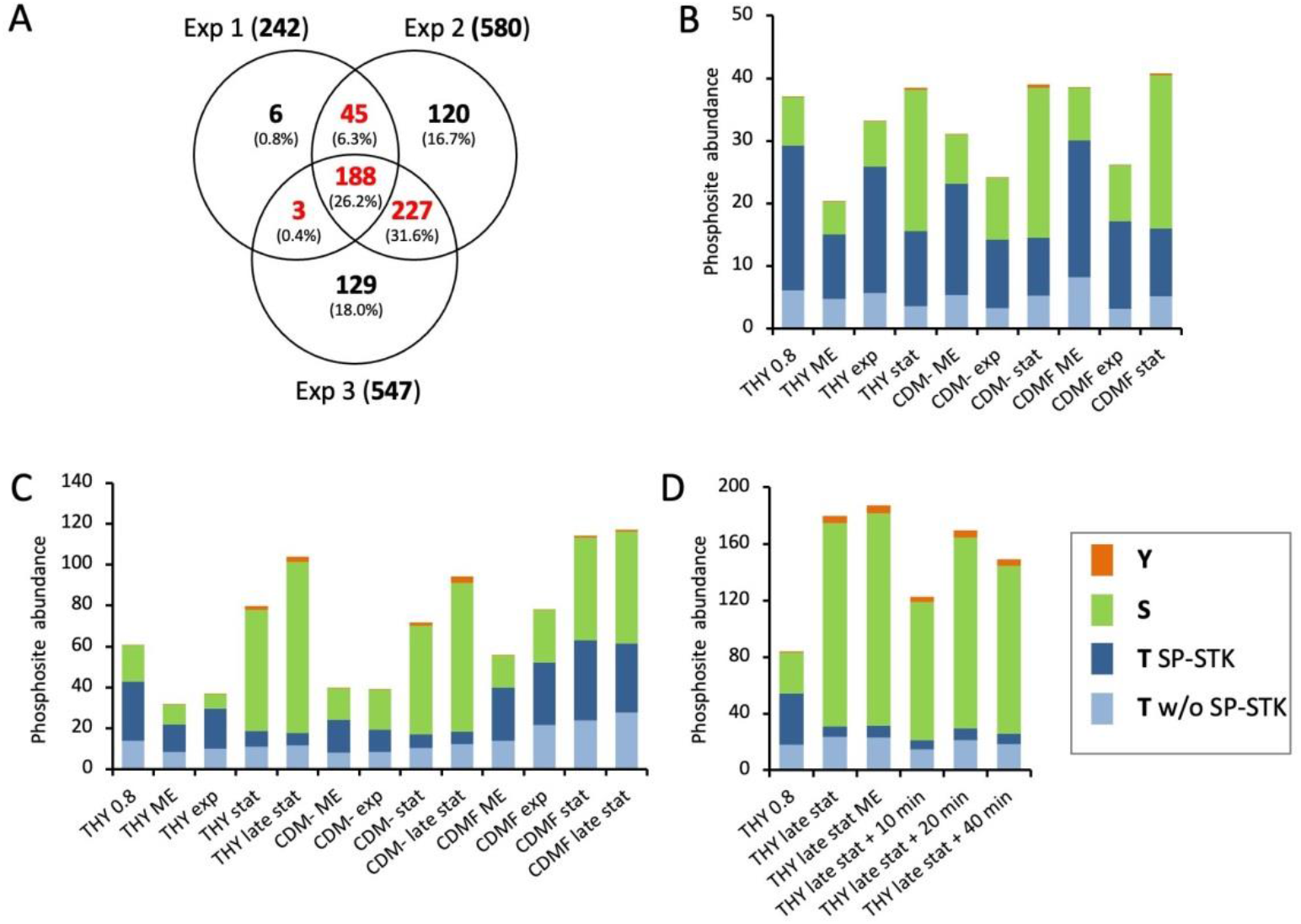
Quantitative analysis of dynamic protein phosphorylation. **(A)** Venn diagram showing numbers and percentages of phosphorylation sites quantified in HDMS^E^ analyses in three experiments. Colored numbers indicate phosphosites that were quantified in at least two experiments and included in the analysis. **(B-D)** Quantitative distribution of phosphorylation events at serine, threonine, and tyrosine in *S. pyogenes* cultures at different growth phases in different culture media in the first (**B**), second (**C**), and third (**D**) experiment. The height of the bars indicates the determined total phosphorylation. The proportions of threonine, serine, and tyrosine sites are color-coded. Threonine phosphorylation of the PASTA kinase SP-STK (T SP-STK) is shown separately from all other threonine phosphorylations (T w/o SP-STK). Results of single phosphopeptide enrichents from single bacterial cultures are shown.

In the first experiment, the proportion of all threonine phosphorylations halved from almost 80% of the total phosphopeptide amount to about 40% in the stationary phase. At the same time, the proportion of serine phosphorylations increased from 20% to 60%. This trend was similar for all three growth media (Figure 3B). In the second experiment, the cultivation of the bacteria was extended to 72 hours. During cultivation in THY and CDM-, threonine phosphorylation decreased even more than in the first experiment to about one-third in the stationary phase after 24 hours and one-fourth in late stationary phase after 72 hours, while the amount of serine phosphorylation and total protein phosphorylation increased. Interestingly, the absolute amount of serine phosphorylation continued to increase after the bacteria entered the stationary phase or decline phase in THY and CDM-. However, in CDMF, the changes in the ratio between threonine and serine phosphorylation were less pronounced in the second experiment than in the first experiment (Figure 3C). This may be related to the differences in the time course of metabolic adaptation to fructose utilization observed in the analysis of the proteome. The third experiment confirmed the reversal of the ratio of threonine and serine phosphorylations between the initial culture and the late stationary phase in THY. Transfer of cultures from late stationary phase to fresh medium had no significant effect on phosphopeptide abundance during a 40-minute period (Figure 3D).

During growth, 67% to 79% of threonine phosphorylation resided at SP-STK. In particular, T324 of the kinase was by far the most frequent phosphorylation site. The observed decrease in the sum of all threonine phosphorylations towards stationary and decline phases was largely due to the decrease in the phosphorylation level of SP-STK. The sum of all other threonine phosphorylations did not alter significantly, but its composition changed, as will be shown below. Tyrosine phosphorylation also increased in the stationary and decline phases but accounted for only a small quantitative fraction of total phosphorylation. It should be noted that results of single phosphopeptide enrichents from single bacterial cultures are shown, giving a large margin of error for individual time points. However, the described main trends were clearly reproduced by the experiments.

Next, we analyzed the individual phosphorylation events during the course of the growth experiments. To exclude the influence of altered protein expression, phosphopeptide values were normalized against the corresponding protein abundances. Hierarchical clustering was performed to visualize groups of differentially phosphorylated sites (Figure 4, Supplementary Figure S1B). Cultures grown in CDMF were excluded from the cluster analysis due to their inconsistent growth characteristics. Clustering of all growth conditions including CDMF is provided in the supplementary material (Data Sheet 2).

**Figure 4.**
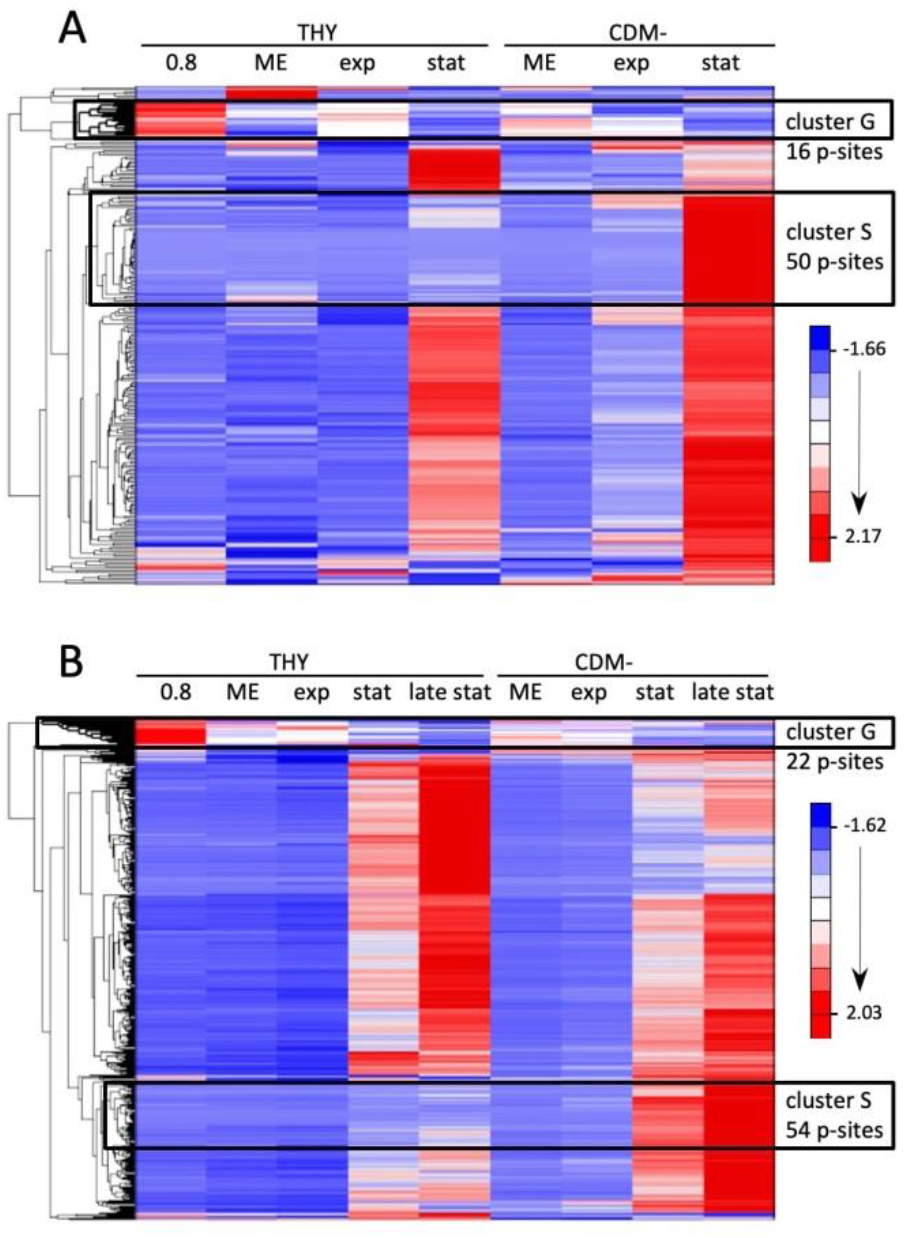
Hierarchical clustering of protein level-normalized phosphorylation site abundances in *S. pyogenes* cultures at different growth phases in THY and CDM-. **(A)** Quantitative data of 236 phosphosites from the first experiment (color coded in Fig. 3A) are included. **(B)** Quantitative data of 460 phosphosites from the second experiment (color coded in Fig. 3A) are included. The G and S clusters are outlined and the number of phosphorylation sites in each is indicated.

The phosphorylation status of most sites increased in the stationary phase. Only one clearly defined cluster (designated as cluster G for growing cells) in the first and second experiment contained sites whose phosphorylation level was highest in the initial cultures and significantly reduced in the stationary or decline phase in THY and CDM- medium (Figure 4). A corresponding cluster was found in the third experiment, in which the initial culture was compared only with late stationary phase cultures (Supplementary Figure S1B). The clusters from the three experiments contained largely consistent phosphorylation sites, 20 of which are listed in Table 2. Notably, 17 out of the 20 phosphorylation events occurred at threonine residues, indicating that threonine phosphorylation is greatly overrepresented in this group. Most of the phosphorylation sites, including four phosphosites of SP-STK, belong to proteins involved in the cell cycle of Gram-positive bacteria. In addition, some peptides whose phosphorylation sites could not be localized with certainty showed comparable phosphorylation profiles with the phosphosites in Table 2. These phosphopeptides derived from the cell division proteins SepF (putative phosphosites T121), FtsA (putative p-site S416 or T417) and FtsZ (putative p-site T336), and from APH domain-containing protein (putative p-site T3), endolytic murein transglycosylase (putative p-site T89) and MacP ortholog Spy49_0377 (putative p-sites T7 und T19) (Supplementary Table S2).

**Table 2.** Phosphosites whose phosphorylation level was highest during growth and decreased in the stationary phase and during starvation.

| UniProt Accession | Protein name | p-site | Exp 1 | Exp 2 | Exp 3 | Birk<sup>1</sup> et al. 2021 | Reference<sup>2</sup> |
| --- | --- | --- | --- | --- | --- | --- | --- |
| A0A0H3BY55 | Non-specific serine/threonine protein kinase | T302 | +<sup>3</sup> | + | + | - |  |
| A0A0H3BY55 | Non-specific serine/threonine protein kinase | T316 | + | + | + | + |  |
| A0A0H3BY55 | Non-specific serine/threonine protein kinase | T324 | + | + | + | + |  |
| A0A0H3BY55 | Non-specific serine/threonine protein kinase | T291 | n.i. | + | + | + | a |
| A0A0H3BZ18 | Cell-division initiation protein | T201 | +<sup>3</sup> | + | + | + | a,b,c,d,e,f |
| A0A0H3BZ18 | Cell-division initiation protein | T245 | +<sup>4</sup> | + | + | + | b |
| B5XMJ7 | Cell cycle protein GpsB | T66 | + | + | + | + | f |
| B5XMJ7 | Cell cycle protein GpsB | T86 | + | + | + | + | f,g,h |
| A0A0H3BZR7 | Mid-cell-anchored protein Z | T11 | + | + | + | + |  |
| A0A0H3BZR7 | Mid-cell-anchored protein Z | T42 | + | + | + | + |  |
| A0A0H3BZ23 | Cell division protein FtsZ | T7 | + | + | n.i. | - | a,e |
| A0A0H3BZC1 | Uncharacterized protein Spy49_0377 | T30 | + | + | + | + | e,i |
| B5XJ02 | UPF0297 protein Spy49_1751c | T7 | +<sup>3</sup> | + | + | + | a,d,e,f,g,h,j,k,l |
| B5XI23 | Protein translocase subunit SecA | T809 | + | + | + | + |  |
| A0A0H3BXH0 | Endolytic murein transglycosylase | T122 | +<sup>3</sup> | + | n.i. | + |  |
| A0A0H3C2P8 | Uncharacterized protein Spy49_1748c | T13 | + | + | + | + |  |
| A0A0H3C0J7 | Phosphocarrier protein HPr | S31 |  | + | + | - | f |
| A0A0H3C2S7 | Uncharacterized protein Spy49_1801c | S148 | + | + | n.i. | + |  |
| A0A0H3C2S7 | Uncharacterized protein Spy49_1801c | T125 | + | + | n.i. | + |  |
| A0A0H3BZP2 | Arsenate reductase | S131 | +<sup>4</sup> | + | + | + |  |

<sup>1</sup> Phosphosite was also identified in *S. pyogenes* M1 (Birk et al. 2021).
<sup>2</sup> References are given for sites identified by mutant analysis as targets of PASTA kinases in other bacteria: a – Ulrych et al., 2021; b – Henry et al., 2019; c – Hirschfeld et al., 2020; d – Zhang et al., 2017; e – Silvestroni et al., 2009; f – Hu et al., 2021; g – Iannetta et al., 2021; h – Kelliher et al., 2021; i – Fenton et al., 2018; j – Prust et al., 2021; k – Wamp et al., 2020; l – Hall et al., 2013.
<sup>3</sup> Phosphosite was not included in cluster G of the first experiment, but was located directly next to it.
<sup>4</sup> Phosphosite clustered separately in the first experiment, although the phosphorylation level decreased at stationary phase.
n.i. not identified

The vast majority of phosphosites showed detectable phosphorylation only in the stationary phase or during starvation. Even in starving bacteria in CDM-, whose optical density decreased considerably throughout the experiment, phosphorylation increased during the prolonged cultivation of 72 hours. The increase was absolute and not just a result of normalization to decreasing protein levels. A particularly striking increase in phosphorylation in CDM- was characteristic of two clusters comprising 50 and 54 phosphorylation sites in experiment 1 and 2, respectively (designated as cluster S for starving cells in Figure 4). Together they contained 71 different phosphosites of which 33 were present in both clusters (Figure 5A). By far the largest proportion of these shared phosphosites originated from ribosomal proteins. Additionally, there was a high proportion of ribosomal phosphorylation sites among the phosphosites unique to each cluster. Thus, of the 50 phosphosites forming cluster S from experiment 1, 36 originated from ribosomal proteins, which corresponds to 84% of all ribosomal phosphosites found in experiment 1. Similarly, 36 ribosomal phosphosites, corresponding to 53% of all ribosomal phosphosites found in experiment 2, were enriched in cluster S of experiment 2 (Figure 5A). The degree of phosphorylation of ribosomal proteins also increased in the stationary phase in THY, but was highest in CDM- (Figure 5B,C). In the proteome, many ribosomal proteins decreased particularly strongly in the extended stationary phase in CDM- compared to the other culture media (Supplementary Table S1). However, they were not all and not uniformly reduced, as described for the prolonged stationary phase *E. coli* (Reier et al., 2022). We therefore suspected a relationship between the stability of individual ribosomal proteins and the degree of their phosphorylation, but could find no evidence for this assumption in our data.

**Figure 5.**
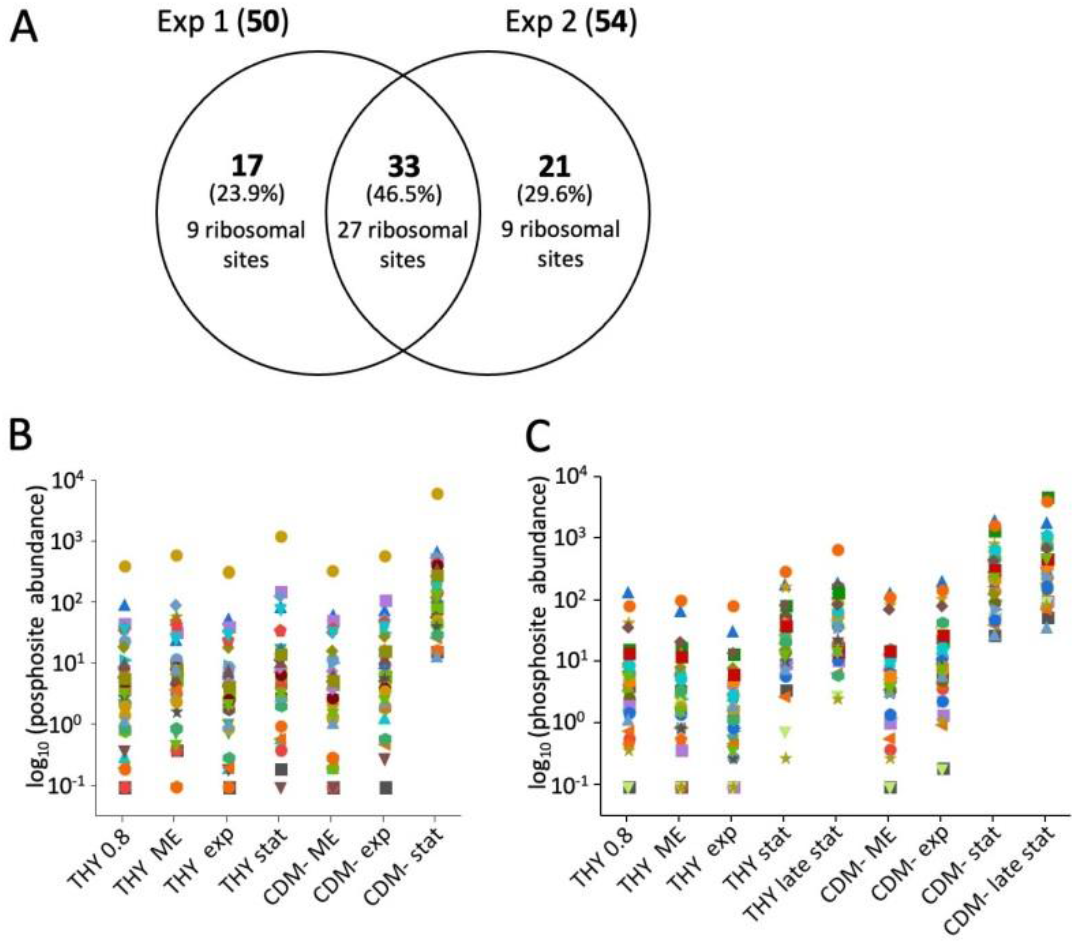
Increased phosphorylation of ribosomal proteins during starvation. **(A)** Venn diagram showing the number and percentage of phosphorylation sites and the number of ribosomal sites in the S cluster of the first and second experiment (see Fig. 3), **(B)** Culture condition-dependent abundance of 36 ribosomal phosphorylation sites included in the S cluster of the first experiment, **(C)** Culture condition-dependent abundance of 36 ribosomal phosphorylation sites included in the S cluster of the second experiment. Phosphorylation site values were normalized against the corresponding protein abundances.

Our quantitative analysis showed that a small fraction of phosphorylation sites, mainly threonine residues, were most phosphorylated during growth, whereas the majority of phosphorylations increased during stationary phase or starvation. Therefore, we wondered if such opposite phosphorylation events also occur within the same protein or peptide. This was indeed the case for many of the proteins listed in Table 2, as exemplified for SP-STK and mid-cell-anchored protein Z in Figure 6. The phosphorylated residues T302 and S299 of SP-STK are located on the same tryptic peptide. During the stationary phase in THY, the phosphorylation of T302 decreased in parallel to the phosphorylation intensity of the very abundant phosphosites T324 and T316, while the phosphorylation of S299 slightly increased. However, in CDM- the phosphorylation intensity of T302 did not decrease indicating nutrient-dependent differences in the dynamics of phosphorylation (Figure 6A,B). The mid-cell-anchored protein Z showed a strong decrease in phosphorylation intensities of T11 and T42 with a concomitant increase in phosphorylation of S2, S5, and S167 during prolonged stationary phase in THY and CDM- (Figure 6C,D). Opposite phosphorylation dynamics of nearby threonine and serine residues located on the same tryptic peptide were also exhibited by the cell cycle protein GpsB (T86/S84), the cell division protein FtsZ (T7/S4), and the protein translocase subunit SecA (T809/S806) (Supplementary Figure S11).

**Figure 6.**
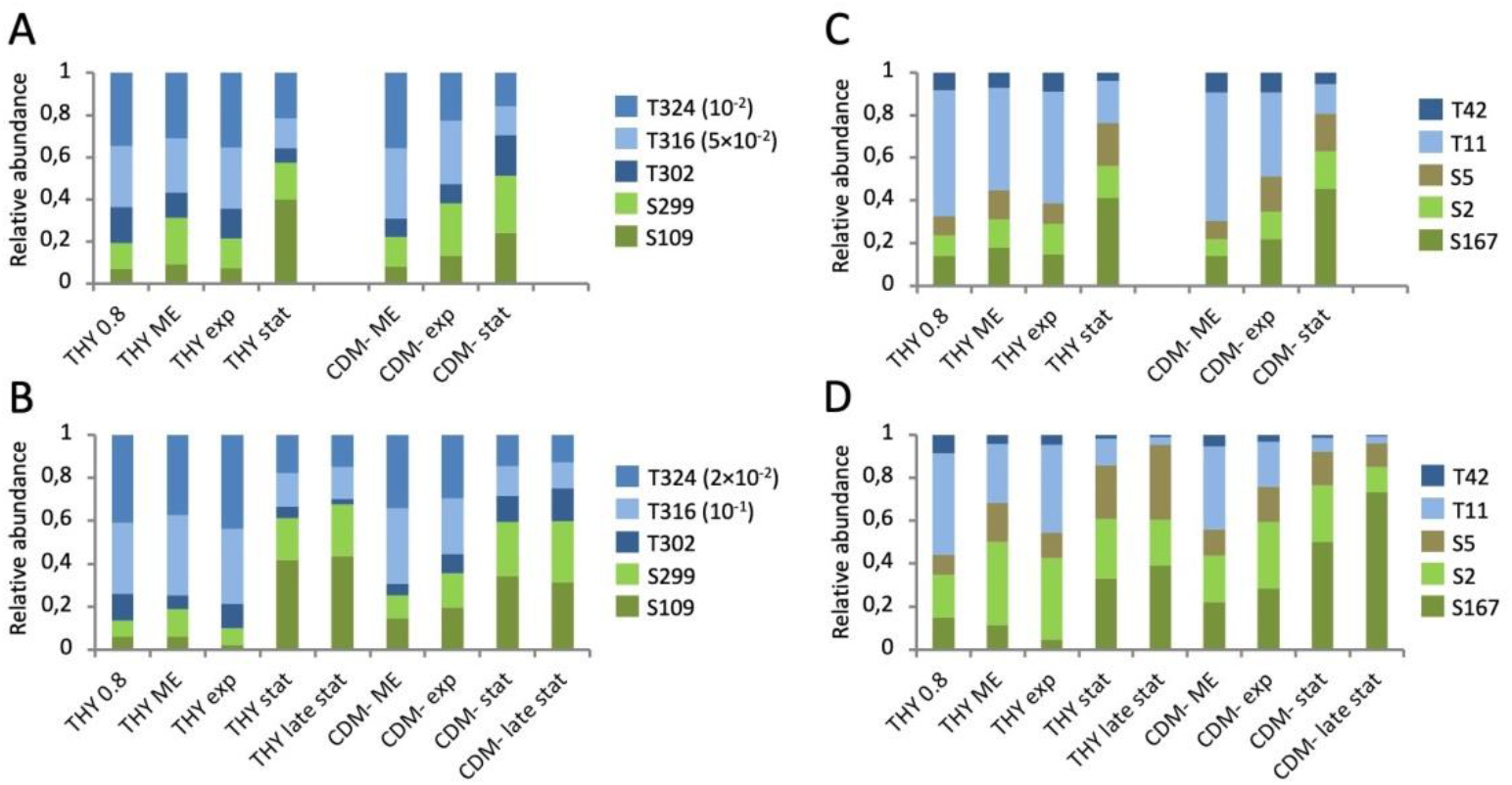
Growth phase-dependent phosphorylation of threonine and serine residues located on the same peptide or protein. **(A,B)** Phosphorylation dynamics of SP-STK in the first **(A)** and second **(B)** experiment. T302 and S299 are located on the same tryptic peptide of SP-STK. **(C,D)** Phosphorylation dynamics of MapZ in the first **(C)** and second **(D)** experiment.

Growth in CDM with fructose also affected specific phosphorylation sites. Despite the differences in adaptation to fructose utilization described above, we found reproducible phosphorylation events in experiments 1 and 2, such as significantly increased phosphorylation of T155 of ribonucleoside-diphosphate reductase and of S13 of uridylate kinase, proteins involved in deoxyribonucleotide biosynthesis and pyrimidine metabolism, respectively (Supplementary Figure S12). While the relationship of these proteins to fructose utilization is not obvious, proteins of the carbohydrate metabolism also showed changed phosphorylation patterns. Notably, enolase phosphorylation sites S2 and S42 were oppositely phosphorylated, with residue S2 being most phosphorylated in THY and CDM- and residue S42 in CDMF (Supplementary Figure S13).

Finally, we calculated the contribution of individual proteins to the phosphoproteome from the sum of their phosphopeptides. In contrast to the quantitative analysis of the phosphorylation sites described above, phosphopeptides with ambiguous phosphosite localization were incorporated and phosphopeptide abundances were not normalized to total protein amounts. In cells from the late exponential growth phase, SP-STK was by far most abundant, followed by DivIVA and phosphoglucosamine mutase (GlmM). Several cell cycle-related putative substrates of SP-STK were found among the 20 most abundant phoshoproteins, along with PTS components, EF-Tu and GAPDH (Supplementary Figure S14). In late stationary phase cells, GlmM, PTS system components, elongation factors Tu and G, as well as chaperones GroEL and DnaK were highly phosphorylated. SP-STK was still among the most abundant phosphoproteins, but its phosphopeptide amount was about fourfold reduced compared to the exponential growth phase (Supplementary Figure S15). Although GlmM was the most abundant phosphoprotein in the late stationary phase, the degree of its phosphorylation increased only 1.4-fold compared with the exponential phase. GlmM was phosphorylated at a single serine residue universally conserved in the active site of phosphosugar mutases and known for its ability to autophosphorylate (Jolly et al., 2000; Schmidl et al., 2010).

## 4 Discussion

### 4.1 Protein phosphorylation in S. pyogenes M49

We investigated S/T/Y phosphorylation events in *S. pyogenes* M49 at different growth phases in three culture media and identified 955 phosphorylated peptides derived from 352 proteins out of which we extracted 815 high-confidence phosphorylation sites in 294 proteins. 162 identical phosphorylation sites were found in the phosphoproteome of *S. pyogenes* M1 (Birk et al., 2021). In *S. pyogenes* M1, threonine was preferentially phosphorylated, accounting for 55% of the phosphorylation sites. This remarkable difference from our results, in which 73% of the phosphorylation events occurred at serine, will be due, at least in part, to the different culture conditions.

The quantitative analysis of dynamic phosphorylation which included a subset of 463 phosphorylation sites for which HDMS^E^ data were available from at least two of the three experiments revealed the following trends: (i) the sum of Ser/Thr/Tyr phosphorylations increases in stationary phase, (ii) the total amount and the relative proportion of threonine phosphorylation decreases in the stationary phase, primarily due to the decrease of threonine phosphorylation of SP-STK, whose expression does not change significantly in the whole proteome, (iii) the total amount and relative proportion of serine phosphorylation increases in the stationary phase, and (iiii) tyrosine phosphorylation behaves similarly to serine phosphorylation, but its proportion is small (max. 3,4%). Starved bacteria in CDM- exhibited similar phosphorylation patterns as cultures in stationary phase.

### 4.2 Specific threonine residues of cell cycle-related proteins are putative targets of the PASTA kinase SP-STK during exponential growth phase

Hierarchical clustering of phosphosites normalized to protein abundance revealed a clearly differentiated cluster of decreasing phosphorylation events in the stationary phase with largely consistent phosphosites in each of the three experiments. This cluster contains several abundant pT sites including four phosphorylated threonine residues within the juxtamembrane region of SP-STK, of which T324 displayed the highest signal of all identified phosphopeptides. Phosphorylated threonine residues in the juxtamembrane region have been reported in *S. pneumoniae* (Ulrych et al., 2021) and other bacteria and are proposed to contribute to the activation of PASTA kinases and/or the docking of their targets (Manuse et al., 2016). PASTA kinases autophosphorylate threonine residues of their activation loop in response to specific stimuli (Manuse et al., 2016). The corresponding phosphosites were not found perhaps due to their location on a large peptide of 34 amino acids in length which is in the upper size rage of identifiable peptides in our analyses.

The cluster analysis revealed 16 phosphosites derived from 12 proteins with comparable phosphorylation patterns to the pT-sites in SP-STK. Many of these proteins belong to the divisome, a protein complex assembled during cell division (Briggs et al., 2021), and are known to be substrates of the PASTA kinase during regulation of cell division and morphogenesis in *S. pneumoniae* and other Gram-positive bacteria (Grangeasse, 2016). Thus, we wondered if these phosphosites are direct or indirect targets of SP-STK in *S. pyogenes*. Corresponding phosphorylation pattern of the kinase IreK and its substrate IreB following growth signals or cell wall stress were also found in *Enterococcus faecalis* (Labbe and Kristich, 2017). Therefore, we searched whether the equivalent phosphosites of related bacteria have been identified as targets of their PASTA kinases in proteomics analyses using kinase mutants.

The cell-division initiation protein **DivIVA** interacts with several cell division proteins and is known as a prominent substrate of streptococcal StkP (Hammond et al., 2019). We detected its phosphorylation at T201 and T245. PASTA kinase-dependent phosphorylation of T201 is highly conserved among streptococci, having been identified using kinase mutants in *S. pneumoniae* (Hirschfeld et al., 2020; Ulrych et al., 2021), *S. thermophilus* (Henry et al., 2019), *S. suis* (Zhang et al., 2017; Hu et al., 2021), and *S. agalactiae* (Silvestroni et al. 2009). T245 was concordantly phosphorylated in *S. thermophilus* (Henry et al., 2019).

**GpsB** is a DivIVA-like-domain-containing protein present only in Firmicutes (Hammond et al., 2019). In *S. pneumoniae*, GpsB is involved in the coordination of septal and peripheral peptidoglycan synthesis and in the regulation of the levels of StkP-mediated protein phosphorylation (Fleurie et al., PLOS-Genet 2014; Rued et al., 2017; Briggs et al., 2021). GpsB proteins comprise conserved N-terminal and C-terminal domains connected by a poorly conserved linker. The N-terminal domain interacts with the cytoplasmatic membrane (Halbedel and Lewis, 2019) and with penicillin binding proteins (Cleverley et al., 2019) whereas the C-terminal domain provides hexamerization of GpsB (Cleverley et al., 2016). We identified growth phase-dependent phosphorylation of GpsB at T66 and T86. Both sites were also phosphorylated by the PASTA kinase in *S. suis* (Hu et al., 2021). Whereas T66 is located within the less conserved linker region of GpsB, T86 is the first amino acid of the conserved C-terminal domain (Supplementary Figure S16). In *E. faecalis* the orthologous residue T84 was also a substrate of the PASTA kinase IreK (Ianetta et al., 2021). In *S. pneumoniae*, T79, located in the aligned sequences four residues upstream of T86 of *S. pyogenes*, was found to be phosphorylated by StkP (Ulrych el al., 2021). The *B. subtilis* kinase PrkC phosphorylated GpsB *in vitro* at T75, which is located one residue upstream of the conserved threonine residue in the aligned sequences. Unphosphorylated GpsB stimulated autophosphorylation of PrkC, whereas phosphorylation of GpsB reduced the kinase activity, thus providing a negative feedback loop (Pompeo et al., 2015). Although the function of GpsB differs between bacterial species and strains (Rued et al., 2017; Hammond et al., 2019), PASTA kinase-dependent phosphorylation at or near the conserved threonine residue at the beginning of the C-teminal domain appears to be conserved in firmicutes and may serve an important regulatory function in the cell cycle.

Mid-cell-anchored protein Z (**MapZ/LocZ**) is a single-pass membrane protein that locates at the division site and is crucial for correctly placing the FtsZ-ring (Fleurie et al., Nature 2014; Holeckova et al., 2015; van Raaphorst et al., 2017). It was enhanced phosphorylated at T11 and T42 during growth in *S. pyogenes*. MapZ is phosphorylated by *S. pneumoniae* StkP at T67 and T78 (Fleurie et al., Nature 2014; Holeckova et al., 2015; Ulrych et al., 2021; Hirschfeld et al., 2020). Both sites are not highly conserved among the MapZ-possessing streptococci, lactococci, and enterococci (Holeckova et al., 2015), only T78 has an orthologous site (T73) in *S. pyogenes*. Indeed, we identified phosphorylation of T73, but only in the second experiment, so this site was excluded from the quantitative analysis.

The tubulin-like protein **FtsZ** was phosphorylated at T7. Orthologous target sites of the respective kinases were identified in *S. agalactiae* (Silvestroni et al., 2009) and *S. pneumoniae* (Ulrych et al., 2021).

The uncharacterized protein **Spy49_0377** is orthologous to MacP, which was recently characterized as a membrane-anchored cofactor of PBP2a in *S. pneumoniae* (Fenton et al., 2018). MacP is phosphorylated at T32 by StkP and this phosphorylation is required for the activation of the PBP2a peptidoglycan synthase (Fenton et al., 2018). A proteomic study using kinase mutants confirmed the phosphorylation of MacP at T32 in *S. agalactiae* (Silvestroni et al., 2009). We identified the orthologous residue T30 as one of the most abundant phosphorylation events during growth in *S. pyogenes*.

**UPF0297 protein Spy49_1751c** belongs to a group of highly conserved proteins in low-GC Gram-positive bacteria, such as IreB from *E. faecalis* and ReoM from *Listeria monocytogenes*. They have been shown to be substrates of PASTA kinases and to be involved in the control of cephalosporin resistance in *E. faecalis* (Hall et al., 2013) and peptidoglycan biosynthesis in *L. monocytogenes* (Wamp et al., 2020; Kelliher et al., 2021). IreB was phosphorylated *in vitro* at the conserved threonine residues T4 and T7, and T7 was found to be the primary site of phosphorylation (Hall et al., 2013). T7 of IreB/ReoM orthologues was identified as target site of the PASTA kinase in most phosphoproteomic analyses of low-GC Gram-positive bacteria (Table 2). Consistent with these results, we found T7 phosphorylated in *S. pyogenes*. Furthermore, we identified peptides of Spy49_1751c phosphorylated at T4 as well as doubly phosphorylated at T4 and T7.

The peptidoglycan biosynthesis protein endolytic murein transglycosylase **MltG** was phosphorylated at T122. Differing sites of this protein have been identified as targets of PASTA kinases in *S. pneumoniae* (Hirschfeld et al., 2020; Ulrych et al., 2021), *S. thermophilus* (Henry e al., 2019), *S. agalactiea* (Silvestroni et al., 2009), *E. faecalis* (Ianetta et al., 2021), and *S. suis* (Hu et al., 2021) all located within the variable cytoplasmic part of this single-pass transmembrane protein.

Phosphorylation of the secretion motor ATPase **SecA** at T809 also showed comparable phosphorylation dynamics to the pT sites in SP-STK. SecA is a potential target of kinases in *Staphylococcus aureus* (Prust et al., 2021), *S. suis* (Hu et al., 2021) and *Clostridioides difficile* (Garcia-Garcia et al., 2022). Most interesting, in *B. subtilis* SecA is required for membrane targeting of DivIVA (Halbedel et al., 2014), and in *L. monocytogenes* an interaction between DivIVA and SecA2 has been described too (Halbedel et al., 2012). Therefore, it is possible that phosphorylation of SecA at T809 exerts a function in the cell cycle in *S. pyogenes*

Uncharacterized protein **Spy49_1748c** was phosphorylated at T13. This multi-pass membrane protein is a member of the competence-induced protein Ccs4 protein family (IPR016978). A role of PASTA kinases for the regulation of competence induction has been shown for different species of Streptococcus (Echenique et al., 2004; Banu et al., 2010; Knoops et al., 2022).

Out of 12 proteins that possess comparably regulated phosphorylation sites to the pT sites in the juxtamembrane region of SP-STK (Table 2), nine are known or suspected cell cycle-related proteins. These are phosphorylated at sites that, to a large extent, have been shown to be targets of PASTA kinases in related bacteria. Therefore, it is likely that these sites are also phosphorylated by SP-STK in *S. pyogenes*, which would need to be confirmed by further analysis using kinase mutants. Together, our results suggest that the PASTA kinase-dependent cell cycle regulatory processes found in related bacteria are also conserved in *S. pyogenes*.

The above discussed phosphorylation events within the juxtamembrane region of SP-STK and at nine potentially cell cycle-related proteins occurred exclusively at threonine residues. A strong enrichment of phosphothreonine among peptides phosphorylated under the control of PASTA kinases and dephosphorylated under the control of their cognate phosphatases was recently reported for *C. difficile* (Garcia-Garcia et al., 2022) and is obvious from published lists of PASTA kinase substrates of various bacteria (Hirschfeld et al., 2020; Ulrych et al., 2021; Henry et al., 2019; Silvestroni et al., 2009; Ianetta et al., 2021; Hu et al., 2021; Prust et al., 2021). Thus, either the substrate specificities of these kinases are generally biased towards threonine or the regulation of cell cycle events by the PASTA kinases predominantly takes place at threonine while other functions of the kinase may target also serine residues.

### 4.3 Mostly serine residues are increasingly phosphorylated during stationary phase and starvation

Many of the cell cycle-related proteins phosphorylated at threonine residues during growth were phosphorylated at other, mostly serine residues in the stationary phase and during starvation. Some of these opposing phosphorylation events occurred in close proximity on the same peptides. Similar growth phase-dependent phosphorylation of the cell division proteins DivIVA and SepF at threonine and serine residues was found in *C. difficile* (Smits et al., 2022). In our analysis, SP-STK and its cognate protein phosphatase SP-STP were also increasingly phosphorylated at multiple serine residues during the stationary growth phase. It would be attractive to speculate that these phosphorylation events might be specific on-off switches for cell cycle regulation. However, because a large number of proteins with a wide variety of cellular functions exhibit greatly increased serine phosphorylation in the stationary phase, a specific function of serine phosphorylation of cell cycle proteins seems unlikely.

Recently, a sharp increase in the number of phosphopeptides after the onset of the stationary growth phase was observed for *C. difficile* (Smits et al., 2022). Already earlier studies found global increases of protein phosphorylation levels in later phases of growth in *E. coli* (Soares et al., 2013) and *B. subtilis* (Ravikumar et al., 2014). Therefore, the increase in protein phosphorylation after the onset of the stationary growth phase seems to be a general feature in the physiology of bacteria. Accordingly, the doubling of the number of phosphorylation sites in *S. pyogenes* in our work compared with a recently published phosphoproteome of the M1 serotype (Birk et al., 2021) is probably due to the inclusion of the prolonged stationary phase in our analysis.

In accordance with our findings in *S. pyogenes*, components of the translational machinery were increasingly phosphorylated during the stationary growth phase in *E. coli* and *B. subtilis* (Soares et al., 2013; Ravikumar et al., 2014). Phosphorylation of ribosomal proteins influences subunit association, binding sites and translational activity of the ribosomes (Soung et al., 2009; Mikulik et al., 2011). Phosphorylated residues were located mostly solvent accessible on the surface of ribosomal proteins (Soung et al., 2009). In *E. coli*, phosphorylation of bL9 was found to be important for cell survival in starvation stress (Pei et al., 2017).

We identified the largest number of phosphosites within a single protein in EF-Tu. Of the total 20 phosphosites, 14 could be used for the quantitative analysis, all of which increased in abundance during the stationary growth phase. Hyperphosphorylation of EF-Tu in the stationary phase has been observed in studies with various bacteria, for example, 32 phosphosites were recently identified in *C. difficile* (Smits et al., 2022). EF-Tu catalyzes the binding of aminoacyl-tRNA to the acceptor site of the ribosome, but additionally to this canonical function it may fulfill diverse moonlighting functions on the cell surface of bacteria (Harvey et al., 2019). Phosphorylation of *E. coli* EF-Tu at T382, which was first reported in 1993 (Lippmann et al., 1993) and at other threonine residues has been shown to inhibit protein synthesis in Gram-negative and Gram-positive bacteria (Sajid et al., 2011; Castro-Roa et al., 2013; Pereira et al., 2015; Talavera et al., 2018). Among several models for the function of EF-Tu phosphorylation in inhibiting protein synthesis, one describes phosphorylation at *E. coli* T382 or at the equivalent site of other bacteria as a switch that interrupts the conformational cycle and traps EF-Tu in an open state. As a result, sites buried in the closed conformation become accessible to the solvent and can be phosphorylated, leading to hyperphosphorylation (Talavera et al., 2018). This model used phosphosites previously identified in *E. coli* (Macek et al., 2008). Interestingly, we found that the corresponding phosphosites in *S. pyogenes* are increasingly phosphorylated in stationary phase and during starvation. This may indicate their function in regulating translation. On the other hand, the phosphorylation events at the stationary phase occurred at substoichiometric levels as can be deduced from the abundance profiles of the corresponding unphosphorylated peptides. This calls into question their physiological relevance. However, in long-term stationary phase cultures of *S. pyogenes,* most cells lose their cultivability during the first days; only 0.1%-0.01% remained viable for weeks or even much longer depending on the culture medium, type of the growth limiting factor, and pH value (Trainor et al., 1999; Wood et al., 2005; Wood et al., 2009; Savic et al., 2012). Therefore, especially after a prolonged stationary phase, the proteome will comprise proteins from cells with different metabolic activity, only a few of which may become persisters (Wood et al., 2009). It is conceivable that the phosphorylation events in the stationary growth phase of *S. pyogenes* are concentrated to a small subpopulation of surviving cells with high phosphorylation status of their proteins. The importance of phosphoregulation for bacterial quiescence was recently described for *S. aureus* (Huemer et al., 2023). Hence, further work is needed to elucidate the function(s) of enhanced protein phosphorylation in stationary phase and starving cells. It is also unknown which kinases are responsible for these phosphorylation events.

### 4.4 Most proteins can probably be phosphorylated during stationary phase and starvation - is there an unknown, rather non-specific kinase?

The massive increase of phosphorylation during stationary phase was not limited to certain groups of proteins, such as components of the translational machinery. Our finding of phosphopeptides from 91 of the one hundred most abundant proteins indicates a strong correlation between protein abundance and the identifiability of phosphopeptides. It can be assumed that, with the exception of certain proteins whose localization or specific structure prevents their phosphorylation, the vast majority of proteins can be phosphorylated. However, identification of their phosphopeptides will depend on the sensitivity of the proteomic methods used. As previously suspected for different bacteria (Prust et al., 2021; Garcia-Garcia et al., 2022; Huemer et al., 2022), other non-Hanks-type kinases besides PASTA kinase must be involved to phosphorylate the broad substrate spectrum in *S. pyogenes*.

In contrast to many bacteria including *S. pneumoniae*, *S. pyogenes* has no bacterial-type tyrosine kinase (BY-kinase) (Grangeasse et al., 2007). However, it possesses a low molecular weight protein tyrosine phosphatase (Kant et al., 2015) and a member of the recently described class of ubiquitous bacterial kinases (Ubk) (Nguyen et al., 2017). *In vitro*, recombinant UbK of *S. pyogenes* strain M1T1 5448 autophosphorylated on two tyrosine residues and phosphorylated the response regulators CovR and WalR, the Ser/Thr phosphatase SP-STP, and GAPDH, each at several serine and/or threonine and/or tyrosine residues (Kant and Pancholi, 2021). Interestingly, several phosphorylation sites in CsrR (CovR ortholog), SP-STP, and GAPDH, respectively, coincide with our analysis. Of note, recombinant SP-STP und GAPDH used for *in vitro* phosphorylation were slightly phosphorylated already during production in *E. coli* (Kant and Pancholi, 2021) indicating that phosphorylation of the respective sites does not require a very specific kinase. Thus, UbK is an interesting candidate, but to what extend the kinase contributes to phosphorylation in *S. pyogenes*, requires further experiments.

## 5 Conclusions

The quantitative analysis of dynamic protein phosphorylation in *S. pyogenes* during growth, stationary phase, and starvation revealed two main types of phosphorylation events, distinguished by the growth phase in which they predominantly occur and their preference for either threonine or serine. One small group of phosphorylation events occurred nearly exclusively on threonine residues of cell cycle-related proteins and was enhanced in growing cells. Data from the literature support the assumption that these are targets of the PASTA kinase SP-STK. The majority of phosphorylation events occurred in the stationary phase or in starving bacteria. Although there is literature evidence that these phosphorylations may be important for regulatory processes in stationary phase or for persister cell formation, their function and the kinases responsible for their formation need to be elucidated in further analyses. Since our work has shown a crucial influence of growth phase on the phosphoproteome and thus on the activity of kinases, future studies of kinase targets involving kinase mutants should include several different growth phases.

## Author contribution

SM designed the study, performed and evaluated the proteomic experiments, analyzed data and wrote the manuscript. MK analyzed data and performed bioinformatic analyses, NP designed the study, cultivated the bacteria and wrote the manuscript.

## Supplementary material

This article includes Supplementary material.

## Abbreviations

PASTA: penicillin-binding protein and serine/threonine-associated
SP-STK: S. pyogenes-serine/threonine kinase
SP-STP: S. pyogenes-serine/threonine phosphatase
Ubk: ubiquitous bacterial kinase
THY: Todd-Hewitt broth supplemented with yeast extract
CDM-: chemically defined medium without carbon source
CDMF: chemically defined medium with fructose
HDMS^E^: data-independent acquisition mode with ion-mobility separation as an additional dimension of separation
DDA: data-dependent acquisition

